# Sequence Imputation from Low Density Single Nucleotide Polymorphism Panel in a Black Poplar Breeding population

**DOI:** 10.1101/437426

**Authors:** Marie Pégard, Odile Rogier, Aurélie Bérard, Patricia Faivre-Rampant, Marie-Christine Le Paslier, Catherine Bastien, Véronique Jorge, Leopoldo Sánchez

## Abstract

**Background:** Genomic selection accuracy increases with the use of high SNP (single nucleotide polymorphism) coverage. However, such gains in coverage come at high costs, preventing their operational implementation by breeders. Low density panels imputed to higher densities offer a cheaper alternative. Our study is one of the first to explore the imputation in a tree species: black poplar. About 1000 pure-breed Populus nigra trees corresponding to a subsample of the French breeding population were selected and genotyped with a 12K custom Infinium Bead-Chip. Forty-three of those individuals corresponding mostly to nodal trees in the pedigree were fully sequenced (reference), while the remaining majority (target) was imputed from 8K to 1.4 million SNPs using FImpute. Each SNP and individual was evaluated for imputation errors by leave-one-out cross validation in the training sample of 43 sequenced trees. Some summary statistics such as Hardy Weinberg Equilibrium exact test p-value, quality of sequencing, depth of sequencing per site and per individual, minor allele frequency, marker density ratio or SNP information redundancy were calculated. Principal component and Boruta analyses were used on all these parameters to rank the factors affecting the quality of imputation. Additionally, we characterize the impact of the relatedness between reference population and target population.

**Results:** During the imputation process, we used 7,540 SNPs from the chip to impute 1,438,827 SNPs from sequences along the 19 Chromosomes. At the individual level, imputation accuracy was very high with a proportion of SNPs correctly imputed between 0.84 and 0.99. The variation in accuracies was mostly due to differences in relatedness between individuals. At a SNP level, the imputation quality strongly depended on genotyped SNP density and to a lesser extent on the original minor allele frequency. The imputation did not appear to result in an increase of linkage disequilibrium. The genotype densification not only brought a better distribution of markers all along the genome, but also we did not detect any substantial bias in annotation categories.

**Conclusions:** This study shows that it is possible to impute low-density marker panels to whole genome sequence with good accuracy under certain conditions that could be common to many breeding populations.

## Background

In genome-wide analyses, accuracies in association and genomic predictions increase with the density in marker coverage [1, 2]. However, these coverage gains have a high cost, which frequently prevents their operational implementation in breeding programs. Low-density panels imputed to higher densities offer an alternative to systematic genotyping or sequencing. The idea of genotype imputation as supplemental genotyping data was described by Burdick et al. (2006) [3], using the term “in silico” genotyping. In this context, imputation refers to the process of predicting genotyping data not directly available for an individual. Usually, imputation uses a reference panel composed of genotyped individuals with high marker density to predict all missing markers of another panel genotyped at lower density coverage [2]. Imputation can be used in at least three different scenarios: (i) to correct missing data that occurred due to technical problems, (ii) to correct for genotyping errors, and (iii) to infer data for non-genotyped SNPs on a set of individuals [4]. Another more extreme scenario involving imputation is to create all the genotype information of individuals that are no longer available from their extant relatives [5]. Imputation software uses two main strategies: the first is based on pedigree and Mendelian segregation [6–8], and the second relies on linkage disequilibrium [9, 10]. Some authors use sequentially or in a given combination both approaches [11]. The first strategy is the one implemented in algorithms like Lander-Green [12], Elston-Steward [13] or Monte-Carlo sampling algorithms [14, 15]. The second strategy is commonly used for samples with low levels of kinship and unknown ancestors, relying instead on the linkage disequilibrium between markers within the reference population. It uses heuristic algorithms as Expectation Maximization (EM) algorithm, coalescence models and Markov’s hidden strings (HMM) [16, 17]. Recently, a study has compared eight machine learning methods to impute a genotype dataset, but results are of lower quality than those from Beagle, a reference software in the domain of imputation [18, 19] which is based on the forecited second strategy [20]. The imputation accuracy depends on several factors. Among them, there are the genotyping quality, the levels of linkage disequilibrium (LD), the marker density which in turn influences perceived linkage disequilibrium, and the relatedness between reference and imputed populations. Factors affecting imputation accuracy have already been studied both with simulated and empirical data. For instance, Hickey et al. (2012) [21] showed that imputation accuracy increases with marker density. The reference population constitution is also a decisive factor for the imputation accuracy. The reference population should be large enough to capture all relevant haplotypes [6] and recombination events, as well as to estimate correctly LD. The relatedness between the reference and the target panel favours imputation quality, with higher accuracies as relatedness increases between the two groups [22]. The effects of panel size, LD and relatedness become more important with decreasing marker density [6, 23]. Imputation of genotyping data has several advantages, the first being the reduction of genotyping costs [24], which can be very important depending on the species. In addition, imputation of genotyping data also improves the detection of QTLs and the model’s prediction accuracy developed in association studies or genomic selection [2]. The imputation of genotyping data could be used in genetic mapping to enrich genetic maps for a higher coverage. Finally, imputation could correct to a certain degree the eventual heterogeneity in marker density related to constraints in chip design. Such heterogeneity in marker density across the genome happened to be the case of the chip used in our study here [25]. Often, imputation involves a difference in densities between reference and targeted panels of less than 10-fold (i.e. 5K to 50K [26–28] or around 10-fold 50K to 500K [29, 30]). With the increasing access to affordable genomic sequence data, the possibility to use full sequences in the reference panel for imputation becomes a reality, at least for a limited number of individuals. Two studies simulated sequences to find the better strategy between imputation accuracy, number of sequenced individuals and genome coverage [31, 32]. Both studies suggest that a good compromise is sequencing as many individuals as possible but at medium coverage (x8). To our knowledge, only three studies in animals have tried to impute successfully from low and medium densities (13 K and 50-60K) to real sequence data (350K and 13 millions) [33–35]. These studies show that inferring whole sequences from low-density marker panels with good accuracy is possible under certain conditions, notably with high levels of relatedness and persistence of LD between the markers across populations. Our study is one of the first to explore the benefits of imputation to densify SNP genotyping in a forest tree species, usually less favored than livestock in genomic resources. This paper is based on black poplar, specifically on one of the breeding populations that is used to produce hybrid poplars. In the context of this breeding effort, imputation is expected to enrich our knowledge, for the subsequent step of predicting and selecting candidates, in four different aspects: (1) to detect recombination events within chromosomes; (2) to estimate the recombination rate within families to improve subsequent *in silico* predictions of segregation; (3) to enriching the genetic map and (4) to improve genomic evaluation accuracy. The main objective of this study was to demonstrate to what extent high quality imputation was feasible from low density arrays. A complementary objective was to identify the factors that contributed to the quality of the imputation and its impact on the linkage disequilibrium and the annotation profile of covered positions.

## Methods

### Plant material

For this study, 1,039 *Populus nigra* were made available from the French breeding population. This sample was structured into 35 families resulting from 23 parents. Available families resulted from two mating sets. As shown in the **table 1**, the first mating set corresponds to an almost complete factorial mating design involving 4 female and 4 male parents, and resulting in 413 F1 individuals structured into 14 full sib families. The second set involved multiple pair mating schemes involving 8 female and 7 male parents, with a number of crosses per parent ranging from 1 to 5, and resulting in 598 F1 individuals structured into 21 full sib families. Six individuals originated from a collection of French wild populations were also added to the population. All 1,039 individuals in this population were genotyped and 43 of them were also sequenced. Among the sequenced individuals, there were 1 grand-parent, 21 parents, 13 progenies and 2 female individuals that were both progenies in the factorial mating design and subsequently parents in the multiple pair mating set (**table 1**). The progenies to sequence were chosen in such a way that all parents had at least one sequenced offspring. The six sequenced individuals originated from wild populations were added to assess the imputation ability with unrelated individuals. Detail of genotype list and origins were given in **table S1**[see Additional file 1].

**Table 1:**
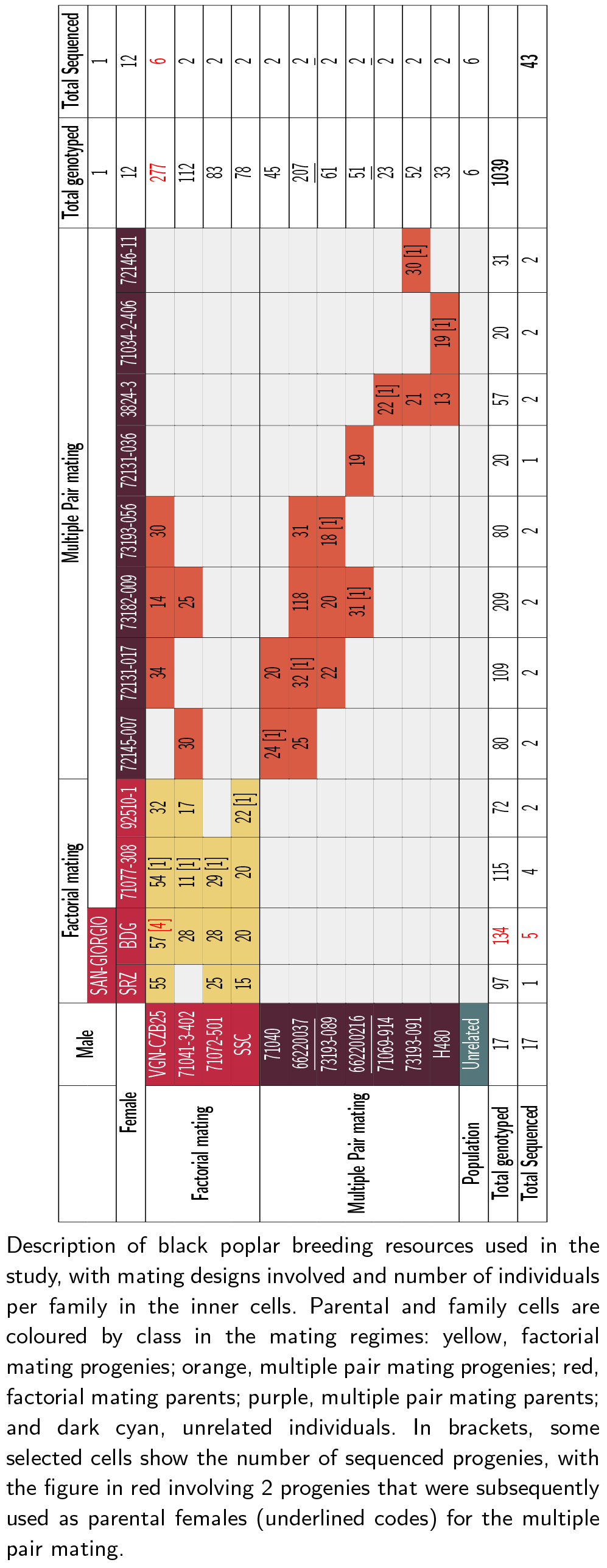
Number of individuals and pedigree information

### Genotyping and sequencing

We used the sequences of 6 parents previously sequenced by Genome Analyzer IIx from Illumina [25]. For the others parents (17), 1 grandparent, 14 progenies and 6 unrelated the DNA extraction was made from leaf samples in the UMR0588-BioForA collection, by using the Macherey-Nagel Nucleospin^®^96 Plant II commercial kit. Illumina paired-end shotgun indexed libraries were prepared from one *μ*g of DNA per accession, using Illumina TruSeq^®^DNA PCR-Free Sample Preparation kit. Briefly, indexed library preparation was performed with DNA fragmentation by AFA (Adaptive Focused Acoustics*™*) technology on Covaris focused-ultrasonicator, all enzymatic steps and clean up were realized according to manufacturer’s instructions. Single or dual indexes were used. Final libraries were quantified by using qPCR using KAPA Library Quantification Kit and Life Technologies QuantStudio*™* Real-Time PCR system. Fragment size distribution of libraries was assessed by High Sensitivity DNA assay either on Agilent 2100 Bioanalyzer or on Caliper LabChip^®^GX nucleic acid analyser. Equimolar pools of multiplexed samples, up to 11, were engaged in sequencing using 4 lanes. After clusters generation on CBot, paired-end sequencing 2 × 150 sequencing by synthesis (SBS) cycles was performed either on a Illumina HiSeq^®^2000/2500 running in high output mode (one lane) or on Illumina HiSeq^®^4000 (three lanes). Reads were trimmed with Trimmomatic (v. 0.32) [36], and mapped to the *P.trichocarpa* version 3.1 genome [37] using BWA-MEM 0.7.12-with default parameters [17]. Picard Tools (v. 2.0.1) [38] were used to remove duplicated reads. Local and Indel realignments were performed using Genome Analysis Toolkit (GATK v. 3.5) [39, 40]. The variant detection was performed on all individuals by two variant callers: (1) in parallel with Freebayes (V1.0.0) [41], and (2) by each individual separately with GATK HaplotypeCaller, to be subsequently assembled using GenotypeGVCFs (called later gVCF-GATK). We have used the VCFtools 0.1.15 [42] to filter variants with no missing data, with a minimum quality score of 30 and a mean depth of 2. We allowed among selected SNPs those harboring three alleles, because mapping was done on another Populus species reference genome, so it was possible to have two alternative alleles and no reference allele in the aligned sequences. We finally kept only SNPs and Indels that were detected by both callers and consistent with Mendelian segregation. To simplify, SNPs and Indels were both called SNPs hereafter. All individuals were genotyped using the Populus nigra 12K custom Infinium Bead-Chip (Illumina, San Diego, CA) [25]. We applied the same quality filters as in Faivre-Rampant et al (2016): markers with more than 90% of missing data were removed and only Mendelian segregation consistent markers were selected.

### Genotype imputation

We used the FImpute software (v 2.2) [11], as many studies have already pinpointed its good performance for imputation when compared to many other alternatives [16, 35, 43, 44]. FImpute can use different sizes of rolling windows with a given overlap to scan the genomes of target and reference datasets. The pedigree information is used to increase imputation accuracy. Therefore, FImpute combines both formerly stated strategies for imputation: that based on pedigree and that on LD. A first round of genotype imputation was performed to predict 1% of missing data still existing on the SNP chip panel. The second and most substantial imputation scheme was between the genotypic data from the chip SNP (SNPchip) and the sequence data (SNPseq). To assess imputation accuracy, a leave-one-out cross validation scheme was performed among the 43 sequenced individuals. The SNPseq were masked for one individual at a time, and this individual with only SNPchip data was subsequently imputed with the rest of individuals. To challenge the imputation scheme, an additional set of 6 unrelated individuals with sequences were added to the target panel. We estimated imputation quality (or accuracy) using various statistics. One was the proportion of alleles correctly imputed by each leave-one-out individual (across SNPs, one proportion per individual and per chromosome: *Propi*), and by positions (across individuals, one proportion per position: *Props*). The proportion of alleles correctly imputed by SNP might be subjected to frequency-dependent bias, in the sense that imputation could be correct more often than not when the imputed allele is already highly frequent. To overcome this, Calus et al. (2014) [45] have proposed the use of an alternative statistic, the Pearson’s correlation coefficient between true and imputed individuals (across SNPs, one correlation value per individual and per chromosome: *Cori*) and between true and imputed positions (across individuals, one value per SNP position: *Cors*). In our case, this latter correlation (*Cors*) was not always available for computation. The reason was that some SNPs had such a low allelic frequency that monomorphic outcomes happened after imputation, leading to zero variances. In order to account for this frequency-dependent outcome, alternatively, we used the option proposed by Badke et al. (2014) [46] to correct the error rate by the probability of correct imputation by chance (*cProps*: corrected SNP proportion). FImpute offers an imputation mode based on allelic frequency (option “random_fill”), which gives us a lower bound for imputation accuracy by individual (*lbPropi* : lower bound individual proportion) and by SNP (*lbProps*: lower bound SNP proportion).

### Factors affecting SNP imputation

We considered different factors describing the heterogeneity between individuals and between markers imputations, and we checked to what extent these factors affected imputation. The first factors were at the individual level: the sequence depth (**MEAN_DEPTH**); and the level of relatedness defined according to the following categories : parent of factorial (Factorial_parents), parent of multiple pair mating design (MultiplePair_parents), progeny of factorial (Factorial_progenies), progeny of multiple pair mating design (MultiplePair_progenies) and French wild population (Unrelated). At SNP level, the following factors were considered: sequencing depth (**DEPTH**) across individuals; per-site SNP quality from the SNP calling step (column **QUAL** in the vcf file, extracted with vcftools v0.1.13 from the gVCF-GATK results files); minor allele frequency (**FreqOri**); the ratio between SNPchip density and SNPseq density in non-overlapping 500kb windows (**RatioDensity**); the p-value of an exact Hardy-Weinberg Equilibrium test (**hweOri**) for each site as defined by Wigginton et al. (2005) [47] and the level of unique information contributed by each SNP given the level of LD with neighbouring SNPs, and calculated as the weight (**Weight**) obtained by the LDAK5 software [48]. The variation of the imputation quality variables (*Props*, *lbProps* and *cProps*) were analysed according to the different factors by a principal component analysis. The factor’s relevance to describe the imputation quality variables were quantified with a Boruta algorithm which is a wrapper built around the random forest classification algorithm implemented in the R (R Core Team 2015) package Borut [49]. This algorithm created “shadowMean”, “shadowMax” and “ShadowMin” attribute values obtained by the shuffling of the original attributes across objects. This set of created attributes is used as a framework of reference. The value of the importance of the factors tested, must be different from the values of the attributes created, to be considered as having importance in explaining the observed variability.

### Linkage Disequilibrium

Plink software [50, 51] was used to estimate the linkage disequilibrium parameter D’ [52] in the SNPchip dataset and after imputation in the SNPseq dataset. These latest were filtered on *Props* (> 0.90) and *cProps* (> 0.80) variables.

### Annotation analysis

We were interested in assessing to what extent imputation could change the annotation profile of covered SNPs, notably given the fact that the process involved a substantial change in density. Changes in annotation profiles from enriched to non-enriched but denser genotypes could be of relevance when using the resulting genotypes to fit prediction models for a large spectrum of traits. To get an annotation profile, a gene annotation analysis was performed. The tool Annovar (v. 2017Jul16) [53] was used with the command “–geneanno -buildver” in the Populus trichocarpa v3.1 gene set.

## Results

### Mapping and genotype calling results

Sequence datasets for every individual were mapped on the P. trichocarpa reference genome v.3.1. In average, 91.7% of reads were mapped, 76.5% were paired and only 2.2% were singletons. The genome coverage was calculated by individual, and it varied between 4X and 52X, with a mean coverage of 13X (**table S1**[see Additional file 1]). A total of 27,475,756 SNPs and Indels were detected by gVCF-GATK, whereas 26,489,941 SNPs were detected by Freebayes (**table 2**). After scoring the SNPs on a quality criterion (Phred score ¿30), the number of trimmed positions were twice as many with gVCF-GATK than with Freebayes (**table 2**). Among the remaining positions, some were monomorphic within *P. nigra* individuals but different from the reference sequence: about 1 million for gVCF-GATK and twice as much for Freebayes. A total of 2,488,736 positions were common between the two callers at that point of the filtering. Among these positions, 17% were Indels and 83% SNPs. To have the best quality in genotype calling, we kept the positions where the genotype calling was at least 95% similar between the two callers for all individuals. Mendelian segregation was checked on available trios, and 142,974 positions were removed for which the progeny were inconsistent with parents. For the chip, after applying quality filters, 7,540 SNPs were recovered for the population under study and were used to impute 1,466,586 SNPs from sequences along the 19 Chromosomes. In other words, we imputed 99% of the data.

**Table 2:**
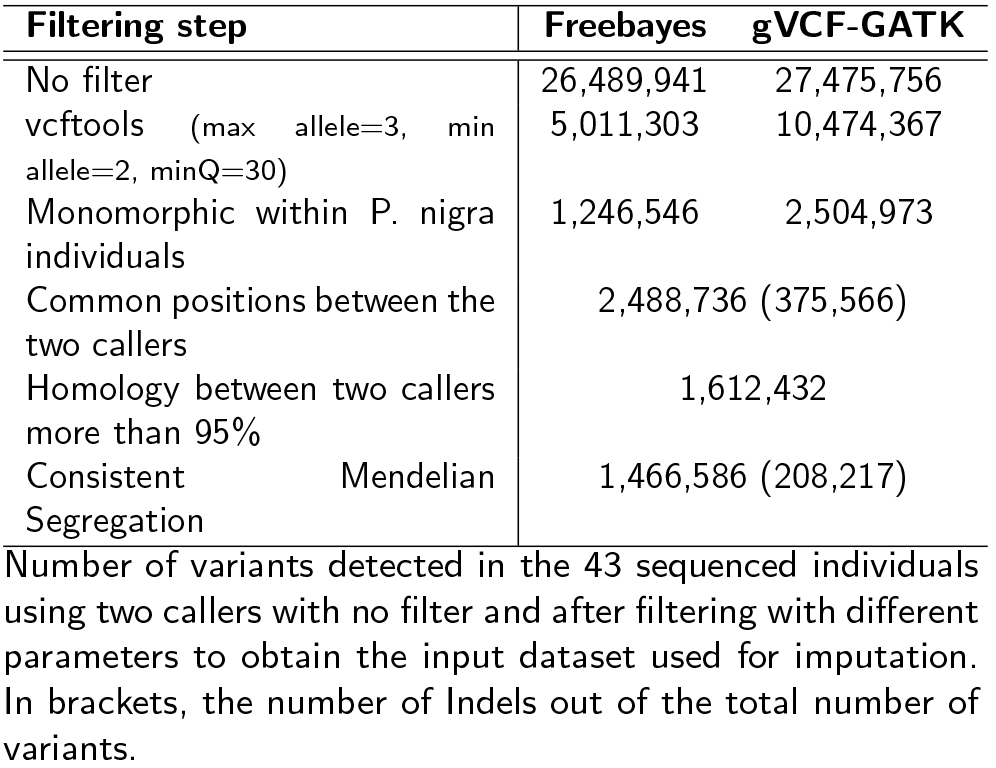
SNP Filter step

### Imputation quality at the individual level

The Pearson’s correlation between true and imputed individuals for each chromosome (*Cori*) was strongly correlated with the individual proportion of SNPs correctly imputed (*Propi*) per chromosome (*R*^2^ = 0.991, **figure 1**), with the former varying between 0.5 and 0.96, and the latter between 0.84 and 0.99. The coefficient of correlation between Cori and Propi was consistently high across individual classes (MultiplePair_parents: 0.929, Factorial_progenies: 0.938, MultiplePair_progenies: 0.929 and Factorial_parents: 0.984), even for unrelated individuals where it was slightly lower with 0.896 (**figure 1**). Propi versus *Cori* relatedness clouds were differently clustered depending on the class of individuals (**figure 1**). In general, factorial mating design progenies had higher *Propi* and *Cori* values (respectively from 0.94 to 0.98 and from 0.81 to 0.95) than those in the Multiple pair mating design progenies (from 0.93 to 0.96 and from 0.80 to 0.88). Progenies from either of the two schemes had higher *Propi* and *Cori* values than those in the parental groups (from 0.87 to 0.90 and from 0.57 to 0.65). The parents of the factorial mating design resulted in the most variable ranges for *Propi* and *Cori* with respectively from 0.88 to 0.99 and 0.6 to 0.96, respectively, although that class had on average higher values than those found in parents in the multiple pair mating scheme. Finally, the unrelated individuals are in the lowest part of *Propi* and *Cori* variation (with respectively from 0.89 to 0.90 and from 0.62 to 0.63). There was no separate group within individual’s categories (**figure 1**) meaning that the individual class ranking was consistent along the chromosomes. The individual lower bound for imputation accuracy (*lbPropi*) was moderately correlated to *Propi* (**figure S1**[see Additional file 2]). The ranking of individual classes was equivalent between *lbPropi* and *Propi*. However, there appears to be a higher gain in *Propi* with respect to *lbPropi* (i.e., using pedigree and LD versus frequencies) for the multiple pair-mating progenies, factorial progenies and factorial parents than for the multiple pair mating parents and unrelated individuals. In **figure 2**, Propi distribution is shown per chromosome. This averaged imputation accuracy was roughly similar for all chromosomes, except for chromosomes 6 and 8 where means were substantially higher (respectively 0.96, and 0.95). No relationship between the sequencing depth (**MEAN_DEPTH**) and *Propi* was found at individual level whereas a poorly significant correlation seems to be present between depth (**MEAN_DEPTH**) and *lbPropi* and *Cori* (**figure S2** [see Additional file 3]). In summary, at the individual level, imputation accuracy was very high with a proportion of SNP correctly imputed ranging between 0.84 and 0.99. The variation was mostly due to the relatedness between individuals and to a lesser extent to sequencing quality or sequencing depth.

**Figure 1:**
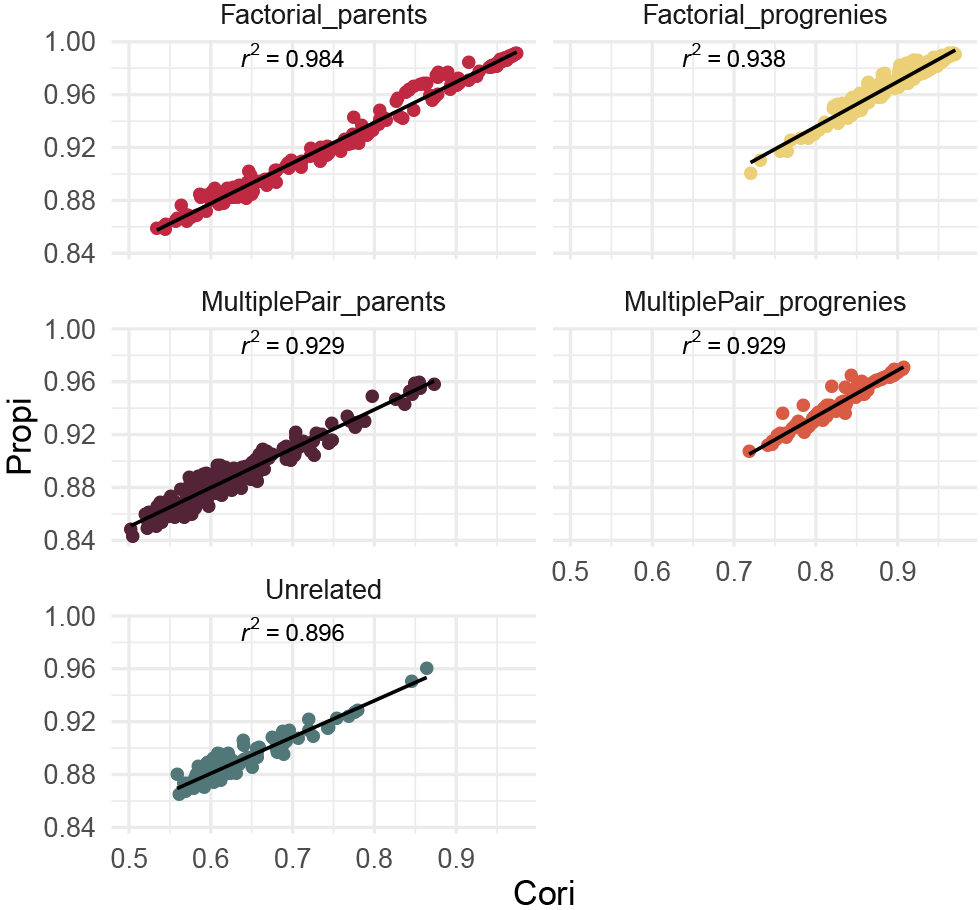
Comparaison of two imputation accuracy variables. Relationship between the proportion of alleles correctly imputed by each leave-one-out individual (*Propi*) and the Pearson’s correlation coefficient between true and imputed individual genotypes (*Cori*). The different panels correspond to the different individual classes in the mating regimes, and each point represents the values for one chromosome and one individual. The correlation value is given in each panel and derives from the fitted regression line.

**Figure 2:**
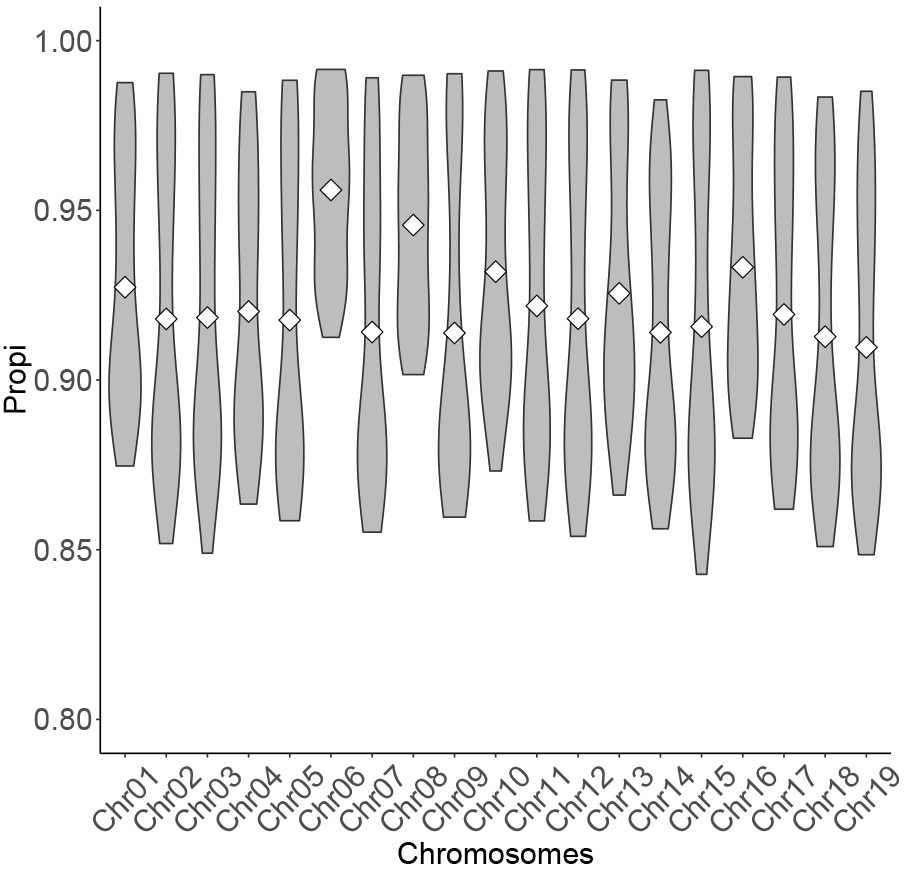
Proportion of individual correctly imputed by chromosomes. Distribution of the proportion of SNPs correctly imputed by chromosomes (*Propi*). White diamond symbol stands for the mean.

### Imputation quality at the SNP level

A very strong correlation between *Cors* and *cProps* (0.94) suggests that similar information was relayed by these two variables despite the frequency-based correction. The **figure S3** [see Additional file 4] shows the variation of the three different estimates of imputation quality at the SNP level (*Props*, *lbProps*, *cProps*), as a function of different classes of minor allele frequency (**FreqOri**). While for low **FreqOri**, *Props* and *lbProps* distributions remained similar, with increasing frequencies their respective distributions tended to separate from each other. The frequency dependent correction applied to cProps was strongest at low frequencies, making *cProps* much lower on average than the other two counterparts. With increasing frequency, that correction was weaker with *cProps* getting closer to both *Props* and *lbProps*. This suggests that, while the problem of sensibility to frequencies can be easily overcome, *cProps* shows imputation qualities that can be far lower than what is actually observed. The first 5 axes of the principal component analysis (PCA) considering the three estimates of imputation quality and six factors that potentially affect this quality, explained 90% of the variance (PC1 and PC2, explained respectively 37.8 and 16.5% of the variation; **figure 3A**). *Props* showed the highest independence with respect to the sequence depth (**DEPTH**), the SNP quality (**QUAL**), *cProps*, the ratio between SNPchip density and SNPseq density (**RatioDensity**) and, to lesser extent, to the level of unique information contributed by each SNP (**Weight**). Props was negatively correlated to the **FreqOri** and positively correlated to the p-value of an exact Hardy-Weinberg Equilibrium test (**hweOri**) and to *lbProps*. In **figure 3B**, correlation of each variable to the PCA dimensions are shown. The first dimension was negatively correlated to **FreqOri**(−0.94), and positively correlated to **hweOri**(0.78), *lbProps* (0.92) and *Props* (0.87). Sequencing quality parameter **QUAL** and **DEPTH** are highly correlated to the second dimension (respectively 0.68 and 0.8). **RatioDensity** and *cProps* were correlated to the third and fifth dimensions whereas the **Weight** variable was only strongly correlated to the fourth dimension. The Boruta analysis ranked the importance of the different factors considered to explain the variation in *Props*, *cProps* and *lbProps* variables (**table 3**). All factors were quantified as being of higher importance than those of lower bond references in shadow attributes. **RatioDensity** resulted in the highest importance among all factors for *Props* and *cProps* with effects respectively being 1,351 and 1,182, largely ahead of the rest of factors, with effects ranging between 40 and 115 for *Props*, 33 and 132 for *cProps*. *lbProps* showed a different ranking of factors, dominated by **FreqOri** with the maximum effect among factors, which is expected given the fact that it is based on allele frequency. In summary, the quality of imputation at a SNPs level strongly depended on **RatioDensity** and to a lesser extent on **FreqOri**. By selecting SNP sets on *Props* and *cProps* simultaneously, we obtained 190,392 SNP with good imputation quality (*Props* > 0.90), while their level of polymorphism was not forced towards low allele frequencies (*cProps* > 0.80). The SNPs distribution along the genome after imputation was more homogeneous than what was initially available with the SNPchip (**Figure 4**).

**Figure 3:**
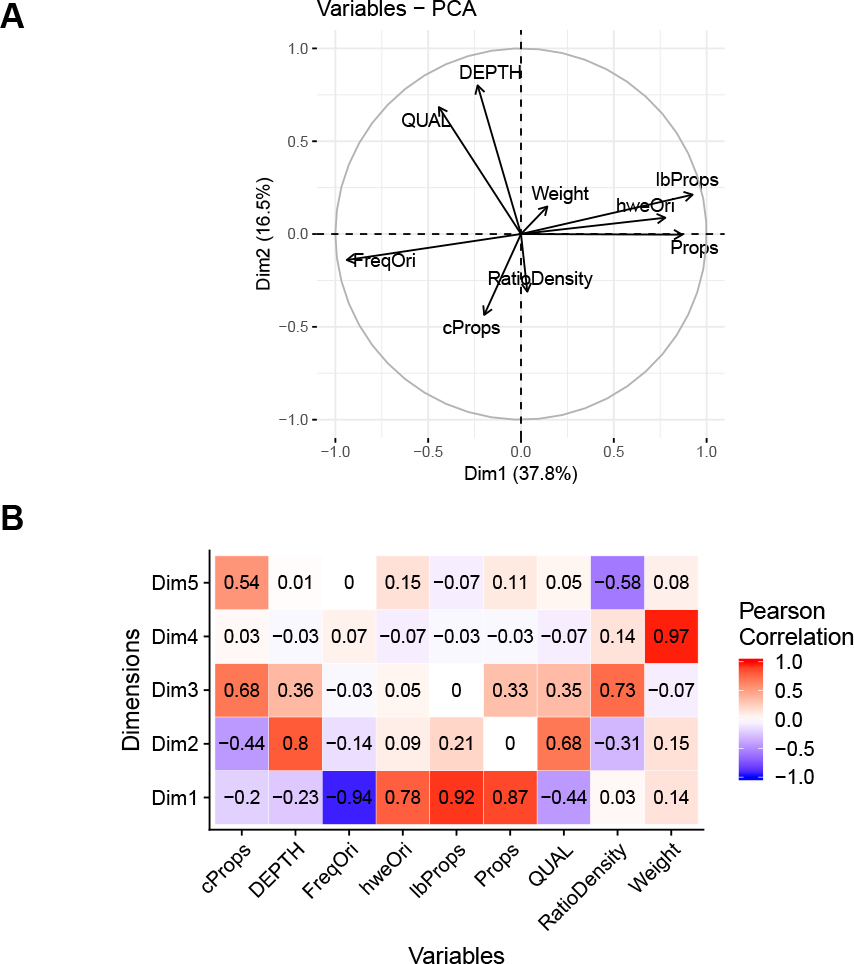
Principal Component Analysis of Factors affecting SNP imputation. (A) Principal Component Analysis factor map of factors calculated at SNP level: *Props*: proportion of SNPs correctly imputed; *cProps*: proportion of SNPs correctly imputed and corrected by the minor allele frequency; *lbProps*: lower bound proportion of SNPs correctly imputed based only on allelic frequency; **hweOri**: p-value of a Hardy-Weinberg Equilibrium test for each site [47];Weight: LD weight estimate obtained with the LDAK5 software; FreqOri: original allelic frequency in the sequenced individuals; **QUAL**: per-site SNP quality from the calling step; **DEPTH**: sequencing depth per site summed across all individuals; **RatioDensity**: ratio between SNPchip density and SNPseq density in a 500kb window. (B) Correlations between parameters calculated at SNP level and dimension of the ACP from figure 3A.

**Figure 4:**
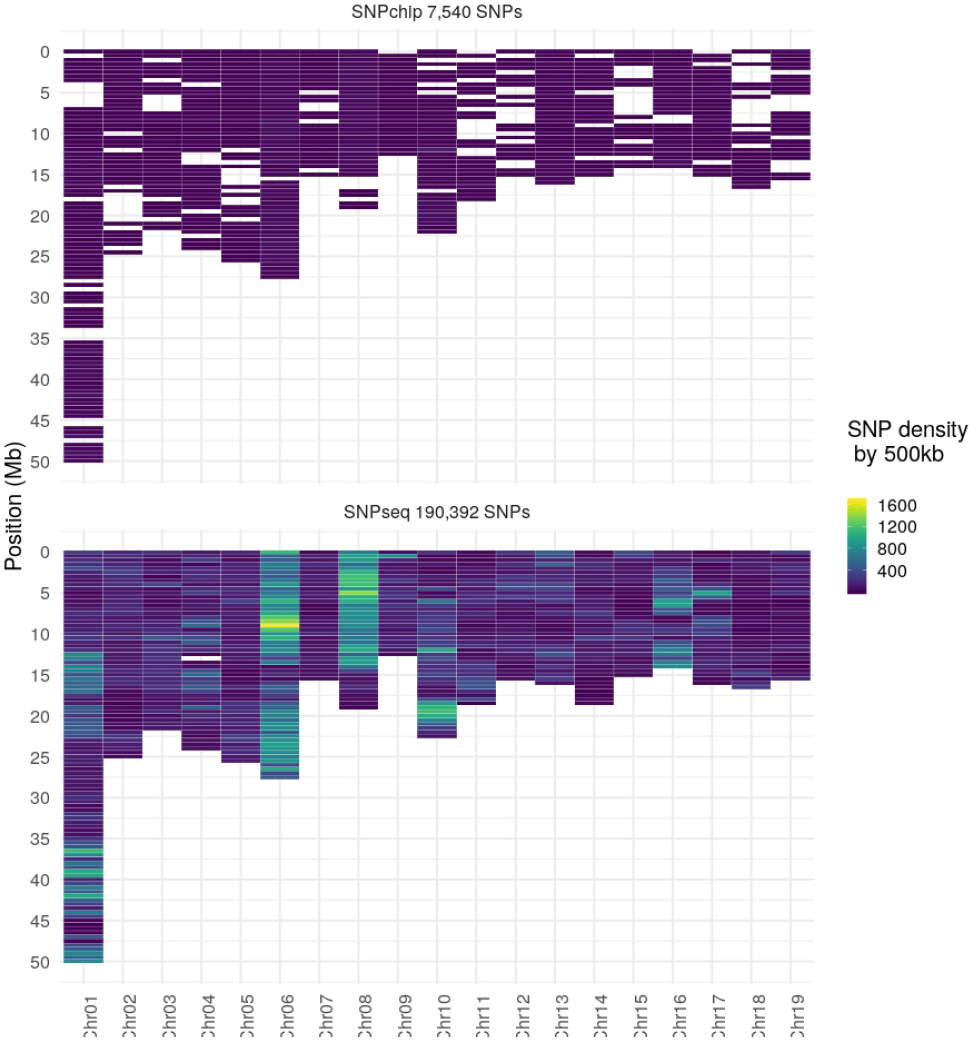
Comparaison of density marker before and after imputation. SNP density map before imputation (top panel), corresponding to the SNP chip genotyping, and after imputation from sequence (bottom) in 500 kb windows. SNPs were selected on two different criteria based on the percentage of alleles correctly imputed: *Props* (> 0:90) and *cProps* (> 0:80). The scale colour represents the density of markers, with dark blue for low density and yellow for high density.

### Linkage Disequilibrium

The linkage disequilibrium (D’) calculated in SNPchip and SNPseq sets is represented in **figure 5A**, with density distributions showing that LD was lower in SNPseq than in SNPchip. This difference between sequence and chip sets was consistent over classes of distances across the genome. **Figure 5B** represents heat-maps for D’ values according to physical distances. In general, D’ decreased with increasing distances, as expected, although this trend was noticeably clearer for SNPchip than for SNPseq. For SNPchip, that D’ decay was noticeable at the very shortest distance lags, with a bottom value for the mean sitting at 0.25. Some increases were observed at the highest distances, but this corresponded to very few number of points. For SNPseq, on the contrary, the weighted mean was almost invariable over distances with a mean value of 0.2. The very large numbers of short distance pairs with low D’ had a high impact on the pattern of the weighted mean. **Figure 5C** presents the results under an alternative view in order to explain the differences in patterns between SNPchip and SNPseq. D’ values are plotted as a function of distance and product of MAF of involved alleles, with the idea of checking to what extent the levels of D’ was the result of low allelic frequencies in SNPseq. For the SNPchip set, the highest values of D’ were found distributed over different distances and levels of MAF products, with a concentration of maximum values at very short distances and relatively low levels of MAF. The picture is substantially different with the SNPseq, where the highest values of D’ were found exclusively at a very narrow band of low frequencies, suggesting that at least part of the levels in D’ could be explained by the low polymorphisms brought by the sequence. As a consequence, the imputation did not appear to result in an increase of LD, but rather the opposite due to the differences in spectra of frequencies between SNPchip and SNPseq.

**Table 3:** Estimation of importance of different explanatory factors by Boruta analysis

| Factor | cProps<br>(Mean ± SD) | lbProps<br>(Mean ± SD) | Props<br>(Mean ± SD) |
| --- | --- | --- | --- |
| shadowMax | 1.44 ± 0.93 | 1.48 ± 0.70 | 1.80 ± 1.30 |
| shadowMean | -0.05 ± 0.79 | -0.22 ± 0.52 | -0.01 ± 0.82 |
| shadowMin | -2 ± 0.53 | -2.22 ± 1.25 | -1.57 ± 0.91 |
| hweOri | 32.96 ± 1.04 | 39.84 ± 0.73 | 40.33 ± 1.99 |
| QUAL | 98.95 ± 5.12 | 67.83 ± 1.70 | 67.90 ± 1.76 |
| Weight | 131.86 ± 3.56 | 92.78 ± 4.20 | 101.57 ± 4.23 |
| FreqOri | 64.28 ± 2.19 | **110.92 ± 2.79** | 115.02 ± 3.28 |
| DEPTH | 114.21 ± 5.08 | 75.51 ± 1.67 | 114.81 ± 4.81 |
| RatioDensity | **1,182.87 ± 39.82** | 36.68 ± 1.50 | **1,351.57 ± 43.94** |

Boruta analyses for the different explanatory factors assumed for imputation quality variables *Props*, *cProps* and *lbProps*. Values correspond to averaged effects and their corresponding standard deviations allowing for a ranking of importance of the factors. The maximum value is bolded. *Props*: proportion of SNPs correctly imputed; *cProps*: proportion of SNPs correctly imputed corrected by the minor allele frequency; *lbProps*: lower bound proportion of SNPs correctly imputed based only on the allelic frequency; **hweOri**: p-value of a Hardy-Weinberg Equilibrium test for each site [47]; **Weight**: LD weight estimate with the LDAK5 software; **FreqOri**: original allelic frequency in the sequenced individuals; **QUAL**: per-site SNP quality from the calling step; **DEPTH**: sequencing depth per site summed across all individuals; **RatioDensity**: ratio between SNPchip density and SNPseq density in a 500kb window. "ShadowMean", "shadowMax" and "ShadowMin" correspond to effects obtained by shuffling the original attributes across objects and used as a reference for deciding which factors are truly important.

### Annotation

A total of 93.4% of SNPchip and 99.79% of SNPseq were annotated (**table 4**). Most categories in the annotation catalog were enriched in the SNPseq compared to the corresponding levels of enrichment in the SNPchip. In general imputation allowed to increase the representation of downstream (from 2.43% to 6.23%), intergenic (from 3.78% to 32.28%), splicing (from 0.04% to 0.1%), and upstream (from 1.8% to 5.71%) regions. On the contrary, the exonic (from 36.18% to 14.96%), intronic (from 37.02% to 30%), UTR3 (from 8% to 6.3%) and UTR5 (from 3.55% to 3.02%) regions were less represented in SNPseq than in SNPchip. Some SNPs were located in or between two regions, there was already the case with SNPchip with upstream;downstream (1.18%) region but other case appeared with the SNPseq as exonic;splicing (0.001%) and UTR5;UTR3 (0.01%). In the exonic region, SNPs were categorized depending on different mutation types. With SNPseq new locations, three new mutation types were represented: frameshift deletion (0.12%), frameshift insertion (0.06%) and non-frameshift deletion (0.04%). Non-frameshift insertion and Stop Loss were enriched (respectively from 0.01% to 0.04%, and from 0.01% to 0.02%), while there were proportionally, less synonymous SNV (from 19.09% to 6.64%) and non-synonymous SNV (from 16.92% to 7.85%), when comparing SNPseq versus SNPchip (**table 4**). No changes were observed, however, for Stop gain (0.15%). In summary, the genotype densification not only brought a better distribution of markers all along the genome, but also no loss in annotation categories.

**Figure 5:**
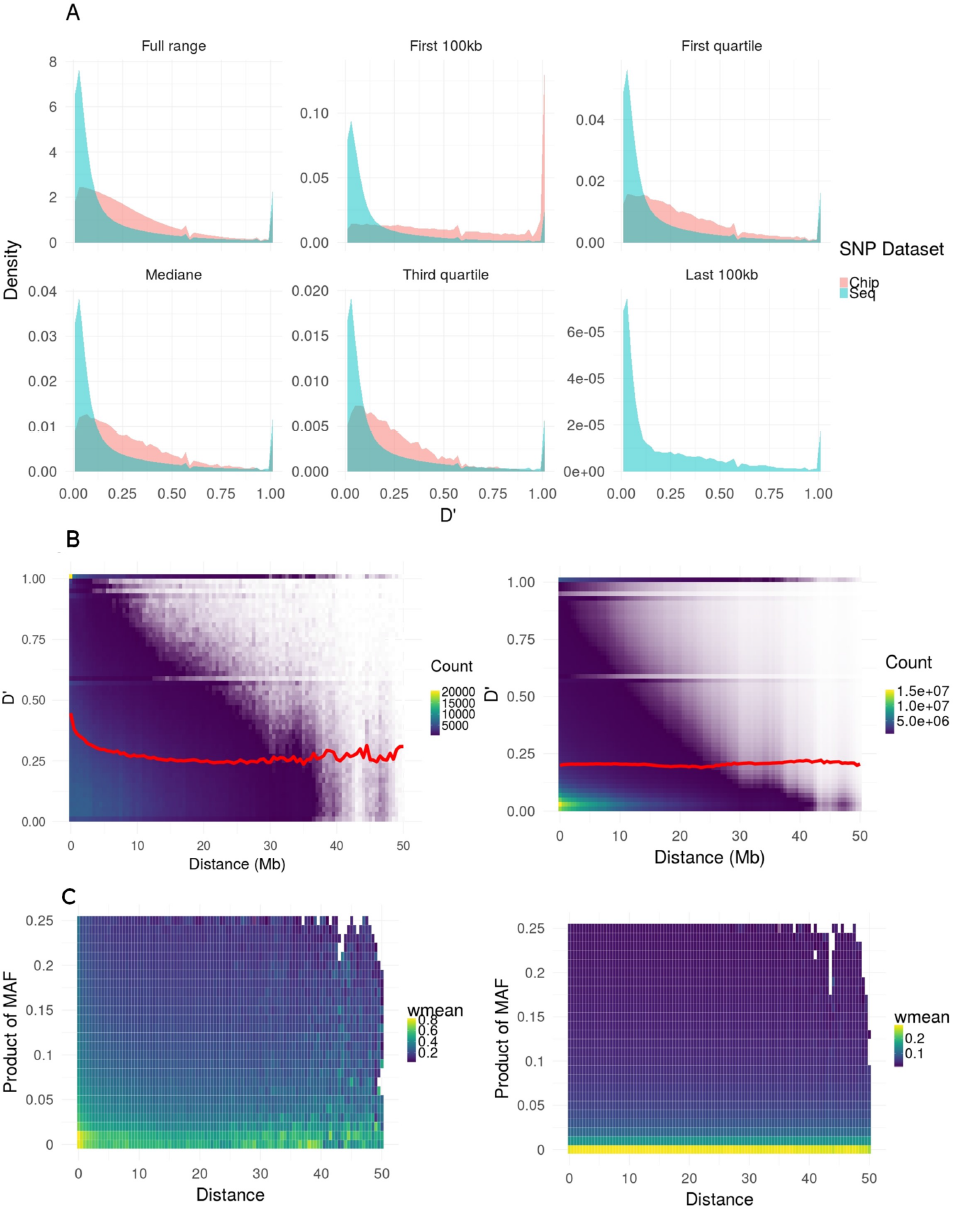
Comparaison of linkage disequilibrium before and after imputation. Distribution of D’ values of linkage disequilibrium for the two SNP sets in the study: SNPchip (pink) and SNPseq (blue) and over different ranges of physical distances (panel A). Panel B represents the distribution of D’ values versus distances in a heat-plot with low densities in blue and high densities in yellow, respectively for SNPchip (left) and SNPseq (right). The red line is the average value of D’ weighted by frequencies for a distance window of 500kb. Panel C represents the distribution of D’ values as a function of distances between any two positions and the product of the corresponding minor allele frequencies in the pair of loci, with colour indicating the average value of D’ weighted by frequencies for a distance window of 500kb from low range (blue) to high range (yellow), respectively for SNPchip (left) and SNPseq (right).

**Table 4:** Proportion of Annotated SNP in genomic regions and mutation types

| Region variant hit | Value % (number) |  |
| --- | --- | --- |
|  | SNPchip | SNPseq |
| downstream | 2,43 (183) | 6,23 (12338) |
| exonic | 36,18 (2728) | 14,96 (29607) |
| intergenic | 3,78 (285) | 32,28 (63901) |
| intronic | 37,02 (2791) | 30 (59377) |
| splicing | 0,04 (3) | 0,1 (192) |
| exonic; splicing | 0 (0) | 0,001 (3) |
| upstream | 1,8 (136) | 5,71 (11307) |
| UTR3 | 8 (603) | 6,3 (12470) |
| UTR5 | 3,55 (268) | 3,02 (5968) |
| UTR5; UTR3 | 0 (0) | 0,01 (20) |
| upstream; downstream | 0,6 (45) | 1,18 (2340) |
| **Annotated Positions** | **93,4 (7042)** | **99,79 (197523)** |
| **Total number of Positions** | **100 (7540)** | **100 (197932)** |
| **Mutation type** |  |  |
| frameshift deletion | 0 (0) | 0,12 (246) |
| frameshift insertion | 0 (0) | 0,06 (118) |
| Non-frameshift deletion | 0 (0) | 0,08 (159) |
| Non-frameshift insertion | 0,01 (1) | 0,04 (84) |
| synonymous SNV | 19,09 (1439) | 6,64 (13133) |
| Non-synonymous SNV | 16,92 (1276) | 7,85 (15534) |
| Stop gain | 0,15 (11) | 0,15 (299) |
| Stop loss | 0,01 (1) | 0,02 (35) |
| **Total number of exonic positions** | **36,18 (2728)** | **14,96 (29608)** |

Annotation results for SNPchip and SNPseq in percentage of counts per annotation category, and number of corresponding positions in brackets. For region variant hit: exonic;splicing corresponds to a variant within exon region but close to exon/intron boundary; UTR5;UTR3 corresponds to a variant positioned where two coding regions overlapped, one in forward and one in reverse; upstream;downstream corresponds to a variant positioned in an intergenic region between two neighbouring genes.

## Discussion

In this study, we have shown that substantial (23-fold) densification in marker coverage is possible in up to 1000 individuals through imputation from a few sequenced nodal individuals (43). Simultaneously, we attained imputation qualities as high as 0.84. This imputation quality is similar to the one obtained on horses [34] with Impute2 software or in cattle [33], and higher than the one obtained on chickens [35]. The study is based on a subset of a breeding population in black poplar, with a relatively low effective number of contributing parents, which could explain partly the success of the imputation. However, this situation is far from exceptional and could be easily found in many other species going through breeding activities, where an elite of a few dozens of parents can contribute substantially to next generation [54]. Although relatedness between the group bringing marker density and the group to be imputed is key in the success of imputation [21, 24, 55], our study demonstrated also that imputation works with relatively small losses in quality when inferring unrelated individuals taken from a diversity collection of the natural range of the species in France. Moreover, such a substantial 23-fold imputation did not appear to increase artefactually the levels of LD. The annotation of imputed positions showed no loss in annotation categories compared to original low density coverage. This two results suggest that imputed data can be of enough quality to be the base of subsequent studies in genome-wide predictions.

The use of a “leave-one-out” cross validation scheme allowed us to ascertain the actual quality of the imputation, both by individuals and by SNP positions. The proportion of alleles correctly imputed by SNP gave the actual value of the imputation quality, although with the drawback of an allele frequency bias. Indeed, a selection based on that proportion by SNP alone could potentially favor positions with low MAF over the rest, as imputation is easier when one of the alternative alleles is rare. The correction we used based on the work of Badke et al. [46] compensated this bias. This measure is interesting whenever we wish to compare results between different imputation methods or between different software. However, it offers a less intuitive criterion, not easily connected to the actual values of imputation error. Therefore, we proposed to combine the actual value of the imputation quality and the frequency-based corrected measure to select SNPs that fulfil both criteria with high level values. Both criteria were given equal importance. The result in our study led to positions with the highest imputation quality while not necessarily resulting in an excess of rare alleles in the imputed population.

Many factors can affect imputation quality like LD, density ratio, minor allele frequency or relatedness between target and reference populations [56, 57]. Our results showed that all these factors considered in our study impacted to various degrees the quality of imputation. It seems difficult to provide general predictor for the imputation quality based on these or other factors. For instance, [4] suggest that there is no obvious pre-imputation filter ensuring a good imputation quality. However, one of the factors with the highest impact on imputation quality in our study was the marker density in the neighborhood of the considered position for imputation. This is a somehow logical outcome, in the sense that numerous markers in dense regions would mutually facilitate their imputation through the extent of LD. These results were consistent with the fact that the imputation accuracy decrease with increasing distance between markers [58]. When designing a low-density chip, it is therefore important to choose SNPs regularly spaced. These results are consistent with the results of He et al. 2018 [59], which showed that an evenly-spaced SNPs combined with an increased minor allele frequencies SNP panel showed the best results.

Imputation requires some degree of LD in existing genomes to reconstruct missing positions [21]. Whenever the reconstruction comprises large chunks of genomes, like in our case here, one could hypothesize that there could be a risk of artefactually increasing the frequency of certain extant haplotypes and, therefore, exacerbate LD among imputed positions. A similar hypothesis has been already proposed by Pimentel et al. [27]. However, what we found appears to be the opposite, with a reduction in D’ from 0.25 in the chip to less than 0.2 in the sequence, on average. The imputed sequence led to D’ values in the low range (close to zero), which could be related to the fact that sequences harbor high number of rare alleles for many positions. Some studies [60, 61] showed that the upper limit of LD between two SNPs is mathematically determined by their difference in MAF. In case of extreme differences, alleles cannot match, even at small distances between SNPs, resulting in low LD. A decrease of LD between SNPs could be problematic for subsequent studies based on imputed data, especially at short distances. Indeed, LD is used to capture the effect of nearby quantitative traits loci (QTL), whenever SNPs are not directly placed on the QTL. This potential loss in capacity to capture QTL effects in the imputed sequences might be compensated for by the genotyping densification, which could extend the reach of markers to unexplored regions involving new QTLs. In summary, genotype densification allowed to have a better repartition of the markers along the genome and in different genomic regions. In our case, the proportion of SNPs in intergenic regions increased with the imputation, this compensated the bias of our low-density SNP chip which was enriched in coding regions [25]. Better marker repartition all along the genome could be useful to detect causal variants, as suggested by Jansen et al. [62]. They showed that with the imputation of missing data, the value of Phred-score genotype quality was improved. This lead to a better genotyping quality, a better causal variant identification in association studies and a better variant annotation. Sequences in our study have brought new spectra of allele frequencies, involving a much higher proportion of rare alleles compared to the chip data, which resulted from a carefully selected set of highly polymorphic markers [25]. While low frequencies could have some interest in diversity studies or kinship assignment [63], their use in the context of genomic evaluation or GWAS would be challenging because of power issues unless the involved rare alleles produce very large effects and are captured with large sample sizes.

From an operational point of view, our results showed that imputation can represent a good strategy to reduce genotyping costs. By using a few well-chosen sequenced individuals in the population, very good imputation results could be obtained and considerably increase the number of SNPs available. It is therefore possible to create a low-density chip to impute at high density via sequenced individuals. This could minimize differences in imputation quality along the genome and avoid any over-representation of certain chromosome regions. This type of strategy can be used in a breeding improvement program on several generations. Yet, it would be required to add high density genotyping or sequences every generation [64] in order to keep a high imputation accuracy. Not doing so could reduce the quality of imputation and result in accumulating errors over subsequent generations. Our study is a first step before using gathered genotypes for genome-wide predictions. The impact of imputation accuracy on genomic selection accuracy was studied by several authors. The genotype densification allowed to increase the genomic evaluation accuracy depending on the architecture of evaluated traits [65, 66]. Moreover, genomic selection accuracy increased with better imputation accuracies [26, 28]. The marker effect estimation could be biased and inbreeding levels could be under-estimated [27], if the imputation accuracy is too low.

## Conclusion

In conclusion, we have demonstrated in this study that high imputation quality is possible even from low density marker sets. The relatedness had an important impact on the imputation quality at the individual level, but it is possible to impute unrelated individuals with a good performance. All factors studied here had an impact on the imputation quality at the SNP level, but there is no obvious way to use their effects as criteria for a pre-imputation filter. The genotype densification towards sequences induced a decrease of linkage disequilibrium, due to the spectra of low allelic frequencies. The densification allowed to correct bias in variant annotation profile of the SNPchip marker set, with a better distribution in all genomic region categories.

## Abbreviations

AFA: Adaptive Focused Acoustics
DNA: DeoxyriboNucleic Acid
EM: Expectation Maximization
F1: first filial generation
GWAS: Genome-wide association study
HMM: Hidden Markov Model
Indels: an Insertion or Deletion of bases in the genome of an organism
LD: Linkage disequilibrium
MAF: Minor Allele Frequency
PCR: Polymerase Chain Reaction
qPCR: quantitative Polymerase Chain Reaction
QTL: Quantitative Trait Loci
SBS: Sequencing By Synthesis
SNP: Single Nucleotide Polymorphism
SNV: single Nucleotide Variation
UTR3: 3’ Untranslated region
UTR5: 5’ Untranslated region

## Declarations

Ethics approval and consent to participate Not applicable

## Consent for publication

Not applicable

## Availability of data

This Whole Genome resequencing project has been submitted to the international repository Sequence Read Archive (SRA) from NCBI RA as BioProject BreedToLast PRJNA483561. The datasets analysed during the current study are available in the INRA Data Portal repository, https://data.inra.fr/privateurl.xhtml?token=0f26535e-4c12-4907-8d2b-ce69e39c1ee0.

## Competing interests

The authors declare that they have no competing interests.

## Funding

This study was funded by the following sources : the INRA AIP Bioressource, EU NovelTree (FP7 - 211868), EU Evoltree (FP6-16322), and INRA SELGEN funding program (project BreedToLast) have funded sequencing and genotyping data. MP PhD grant was jointly funded by INRA SELGEN funding program (BreedToLast) and by Region Centre - Val de Loire funding council.

## Author’s contributions

MP performed the analyses and drafted the manuscript. OR developped the variant calling pipeline scripts and helped for sequence data preparation and bioinformatics. AB, PFR and MCLP provided the sequence and genotyping datasets. CB provided access to plant material as scientist responsible for the Populus nigra breeding program. VJ and LS designed the study, assisted in drafting the manuscript, and obtained funding. All co-authors significantly contributed to the present study. All authors read and approved the final manuscript.

## Acknowledgements

The authors acknowledge The authors acknowledge the ‘GIS peuplier”, the UE GBFOR and PNRGF (ONF) for access, maintenance, and sampling of plant material. The authors want to thank Vincent Segura and Aurélien Chateigner for their valuable discussions; Vanina Guérin and Corrine Buret for their work on DNA extraction for the genotyping and the sequencing; Aurélie Chauveau and Isabelle Le Clainche for libraries construction, sequencing and Infinium genotyping; Elodie Marquand and Aurélie Canaguier for data processing and management. EPGV group acknowledges also CEA-IG/CNG by conducting the DNA QC and by providing access for their Illumina Sequencing and Genotyping platforms.

## Additional Files

Additional file 1 : TableS1.pdf — Sequencing, pedigree and reference information’s of each reference individuals.

Additional file 2 : FigureS1.pdf — Relationship between the proportion of alleles correctly imputed by each leave-one-out individual (Propi) and the lower bound individual proportion of SNP correctly imputed lbPropi). The different colors correspond to the different individual classes in the mating regimes, and each point represents the values for one chromosome and one individual.

Additional file 3 : FigureS2.pdf — Relationship between the sequencing depth and imputation quality variables at individual level. On the top of the diagonal: Pearson’s correlations. The distribution of each variable is shown on the diagonal. On the bottom of the diagonal: the bivariate scatter plots. Additional file 4 : FigureS3.pdf — Variation of the three different estimates of imputation quality at the SNP level (*Props* (Green), *lbProps* (Purple), *cProps* (Orange)), as a function of different classes of minor allele frequency (**FreqOri**).

